# A genome-wide genetic pleiotropy approach identified shared loci between multiple system atrophy and inflammatory bowel disease

**DOI:** 10.1101/751354

**Authors:** Alexey A Shadrin, Sören Mucha, David Ellinghaus, Mary B Makarious, Cornelis Blauwendraat, Ashwin A Sreelatha, Antonio Heras-Garvin, Jinhui Ding, Monia Hammer, Alexandra Foubert-Samier, Wassilios G Meissner, Olivier Rascol, Anne Pavy-Le Traon, Oleksandr Frei, Kevin S O’Connell, Shahram Bahrami, Stefan Schreiber, Wolfgang Lieb, Martina Müller-Nurasyid, Andreas Arnold, Georg Homuth, Carsten O. Schmidt, Markus M. Nöthen, Per Hoffmann, Christian Gieger, European Multiple System Atrophy Study Group, J Raphael Gibbs, Andre Franke, John Hardy, Gregor Wenning, Nadia Stefanova, Thomas Gasser, Andrew Singleton, Henry Houlden, Sonja W Scholz, Ole A. Andreassen, Manu Sharma

**Author notes:** Corresponding author’s contact information: Dr. Manu Sharma PhD, Head, Centre for Genetic Epidemiology, Institute for Clinical Epidemiology and Applied Biometry, University of Tubingen, Tubingen, Germany, Silcherstraße 5 72076 Tübingen, Germany. these authors contributed equally (Sören Mucha, David Ellinghaus). a full list of the European Multiple System Atrophy Study Group members is available in the supplementary material.

## Abstract

We aimed to identify shared genetic background between multiple system atrophy (MSA) and autoimmune diseases by using the conjFDR approach. Our study showed significant genetic overlap between MSA and inflammatory bowel disease and identified *DENND1B, C7*, and *RSP04* loci, which are linked to significant changes in methylation or expression levels of adjacent genes. We obtained evidence of enriched heritability involving immune/digestive categories. Finally, an MSA mouse model showed dysregulation of the *C7* gene in the degenerating midbrain compared to wildtype mice. The results identify novel molecular mechanisms and implicate immune and gut dysfunction in MSA pathophysiology.

## INTRODUCTION

Genome-wide association studies (GWASs) and subsequent GWAS based meta-analysis studies led to the discovery of novel loci for most of the complex diseases [1]. Heritability estimates revealed that studies have explained the fraction of the missing heritability; the majority of the complex diseases are influenced by numerous genes that each have small individual effects [2]. This polygenic architecture of complex diseases necessitated the use of approaches which can leverage the availability of a genome-wide data to identify additional novel loci which can be overlooked by applying a standard genome-wide analytical approach [3]. The evidence generated so far from GWAS has shown overlapping single nucleotide polymorphisms (SNPs) in several phenotypes [4-6]. This genetic-pleiotropy has already been exploited successfully to identify novel loci for various complex diseases, including between Parkinson’s disease (PD) and autoimmune diseases such as Crohn’s disease (CD) and diabetes mellitus type 1 (T1D) [6].

The role of immune dysfunction in neurodegenerative diseases has long been debated. For example, one of the earlier studies using autopsy-confirmed PD cases showed a higher level of microglial activation within the substantia nigra, and this increase in microglia activity was identified by human leukocyte antigen DR (HLA-DR) [7]. With the availability of genome-wide data, GWAS have started to reveal the extent of the immune component in the etiopathogenesis of neurodegenerative diseases [8-10]. Previously published GWAS and follow-up studies in PD provided unequivocal evidence regarding the role of the HLA region in PD pathogenesis [8, 9]. A recently published study identified alpha-synuclein-derived peptides, which regulate the expression of the HLA locus [11]. Likewise, GWAS studies in tauopathies, a group of diseases such as Alzheimer’s disease (AD), frontotemporal dementia (FTD), and progressive supranuclear palsy (PSP) characterized by an abnormal accumulation of neurofibrillary tangles, have established the role of immune component, and thus lending further support to the notion that immune dysfunction is one of the central pathways to neurodegenerative disorders [12-14]. Because of the involvement of alpha-synuclein, multiple system atrophy (MSA) is categorized as a synucleinopathy along with PD, and dementia with Lewy bodies (DLB). MSA is an adult-onset progressive neurodegenerative disorder characterized by a combination of parkinsonism, autonomic, cerebellar or pyramidal signs [15]. Pathologically, it is defined by neuronal cell loss, gliosis and alpha-synuclein positive oligodendroglial cytoplasmic inclusions (GCIs). Neuronal loss in MSA affects striatonigral and olivopontocerebellar structures, and the degree of neuronal loss and GCI density showed a positive correlation between both lesions, suggesting that the accumulation of GCIs is an important factor in neuronal death in MSA [16].

In contrast to other more common neurodegenerative disorders, genetic studies including GWAS have failed to identify disease-associated genetic loci for MSA, though heritability estimates have established a small genetic component in MSA [17]. Nevertheless, emerging evidence points towards deregulation of mitochondria, energy homeostasis, oxidative stress and immune dysfunction that could contribute to the alpha-synuclein inclusions observed in oligodendrocytes in MSA [18]. Despite this, developing a consensus on potential mechanisms that explain the causes of MSA remains challenging. In the present study, we applied a genome-wide genetic-pleiotropy informed approach to identify shared genetic risk factors between MSA and autoimmune diseases with subsequent validation in MSA transgenic mice. Major steps of the study are presented in Figure 1.

**Figure 1.**
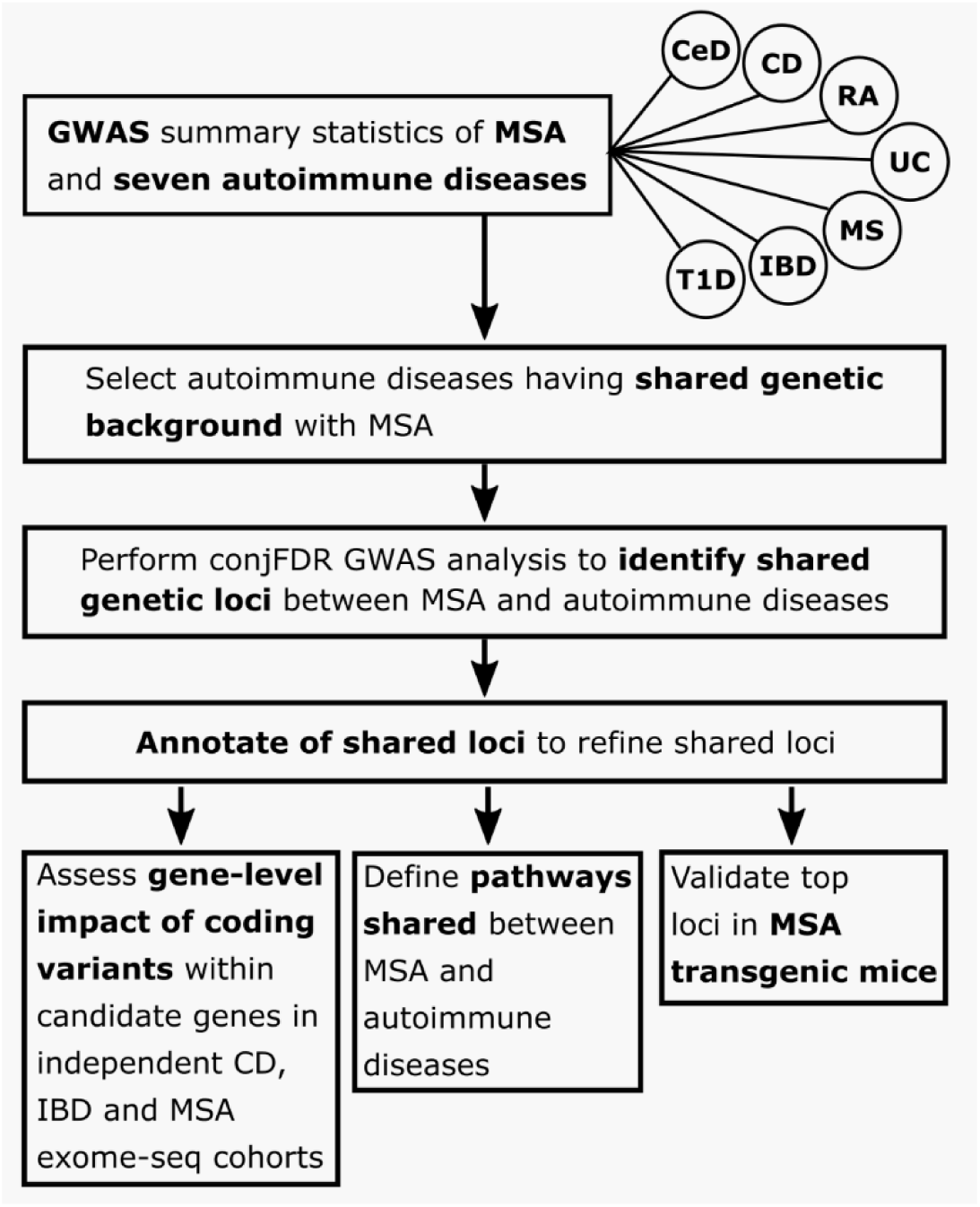
Flow chart representing major steps of the study. MSA: multiple system atrophy; CD: Crohn’s disease; IBD: inflammatory bowel disease, including CD, ulcerative colitis and unclassified IBD cases; UC: ulcerative colitis; T1D: diabetes mellitus type 1; CeD: celiac disease; RA: rheumatoid arthritis; MS: multiple sclerosis.

## RESULTS

### Genetic overlap between MSA and autoimmune diseases

Conditional Q-Q plots for association p-values of MSA and autoimmune diseases showed strong enrichment for Crohn’s disease (CD) and inflammatory bowel disease (IBD, including CD, ulcerative colitis and unclassified IBD cases) (Figure 2 A-D). Successive leftward shifts for strata of SNPs with higher significance in the conditional phenotype indicated that the proportion of MSA-associated SNPs increased considerably with higher levels of association with IBD and CD (and vice versa), suggesting significant shared genetic background between MSA and both IBD and CD (Figure 2). In contrast, there was weak enrichment observed for other analyzed phenotypes (Supplementary Figure 1).

**Figure 2.**
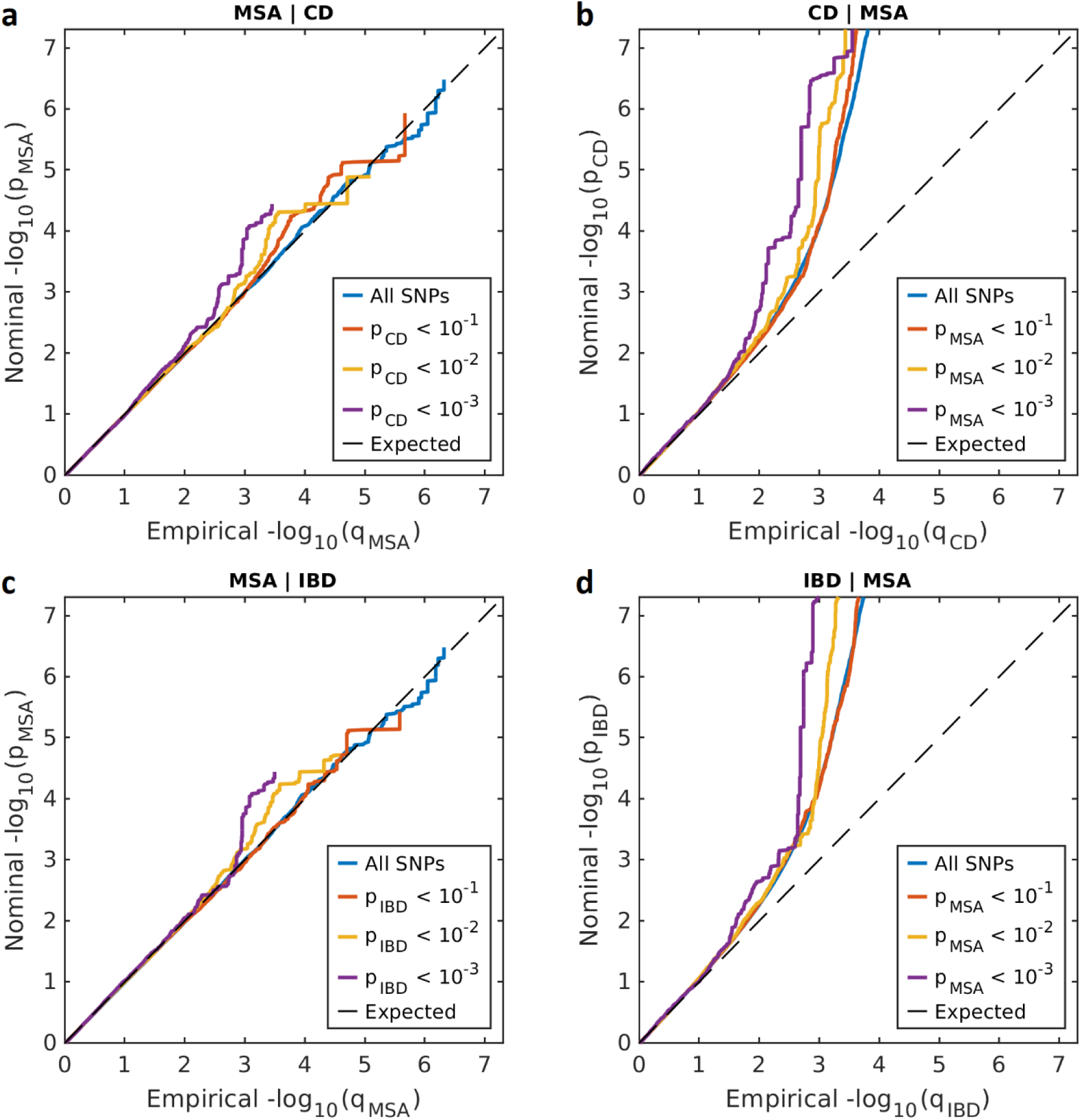
Conditional Q-Q plot showing the relation between expected (x-axis) and observed (y-axis) significance of SNPs in the primary phenotype when markers are stratified by their p-values in the conditional phenotype. A sequence of 4 nested strata is presented: all SNPs (blue), p_conditional_phenotype_ < 0.1 (orange), p_conditional_phenotype_ < 0.01 (yellow) and p_conditional_phenotype_ < 0.001 (purple). Dashed black line demonstrates expected behavior under no association. The increasing degree of leftward deflection from the no-association line for strata of SNPs with higher significance in the conditional phenotype indicates putative polygenic overlap. A: MSA conditioned on CD; B: CD conditioned on MSA; C: MSA conditioned on IBD; D: IBD conditioned on MSA.

### Shared loci between MSA and IBD

Enrichment observed in the conditional Q-Q plot (Figure 2) led us to perform in-depth genome-wide association screening by combining MSA with IBD and CD GWAS summary statistics in conjFDR analysis [19] (see URLs). In the presence of genetic overlap between analyzed phenotypes conjFDR approach offers increased statistical power to discover genetic loci shared between analyzed phenotypes as compared to conventional multiple testing approaches [20, 21]. Three LD-independent regions significantly associated with MSA and IBD/CD were identified at conjFDR < 0.05 (Table 1, Supplementary Table 1). Identified loci with leading SNPs, rs12740041 (*DENND1B*, upstream), rs4957144 (*C7*, intronic) and rs116843836 (*RSPO4*, upstream), in our analysis, suggested underlying common genetic mechanisms between these phenotypes. Manhattan plot for conjFDR results is presented in Figure 3. Detailed regional association plots for identified loci are presented in Figure 4.

**Table 1.**
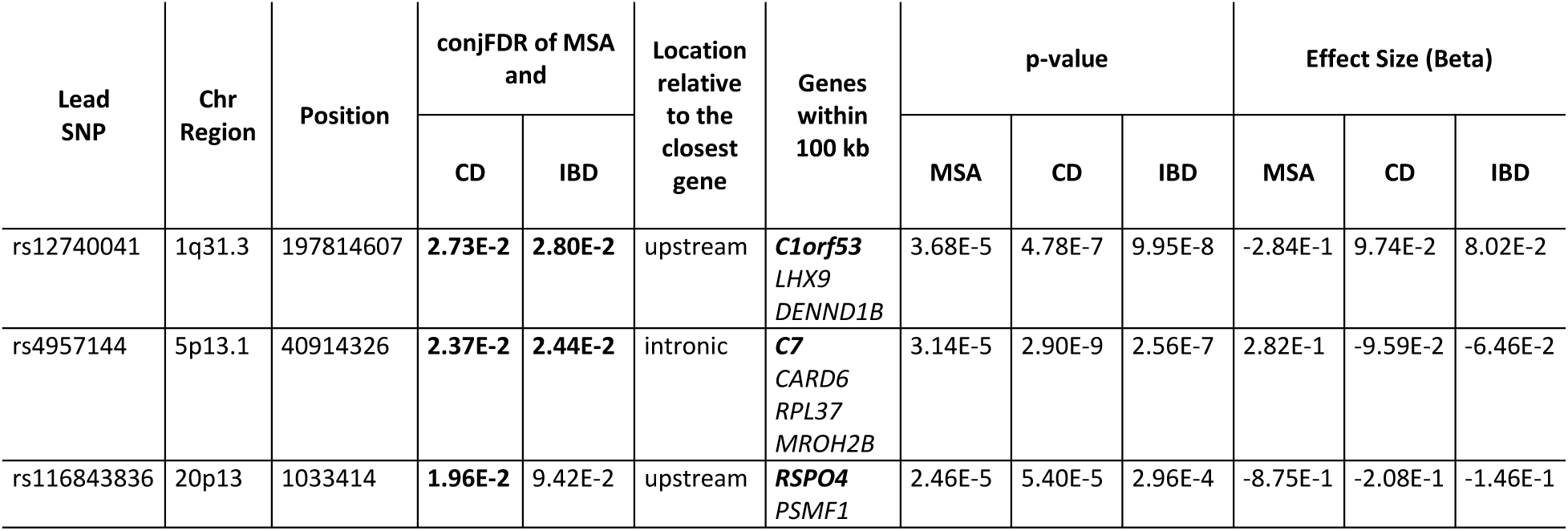
Significant (conjFDR<0.05) loci shared between MSA and IBD/CD. Chromosome (Chr) and position are indicated according to GRCh37. Significant conjFDR values (conjFDR<0.05) are shown in boldface type. Closest gene is shown in bold. For all phenotypes, p-values without genomic inflation correction are shown. The effect size is given as Beta regression coefficient.

**Figure 3.**
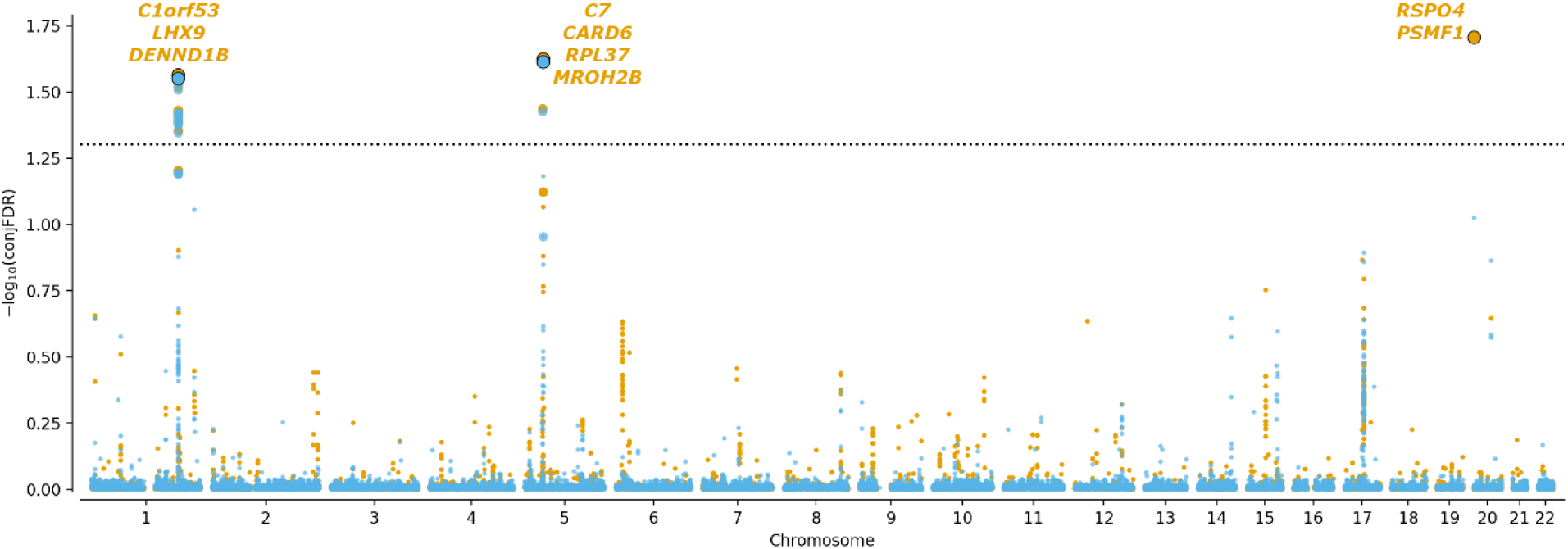
Manhattan plot of -log_10_(conjFDR) for MSA and CD (orange)/IBD (blue). Horizontal dashed black line shows the significance threshold conjFDR=0.05. For each significant locus, genes within 100 kb from the locus lead SNP are shown. Lead variants at each locus are shown as bold dots with black border. Variants in high LD (r^2^>0.6) with the lead variant are shown as bold dots without border.

**Figure 4.**
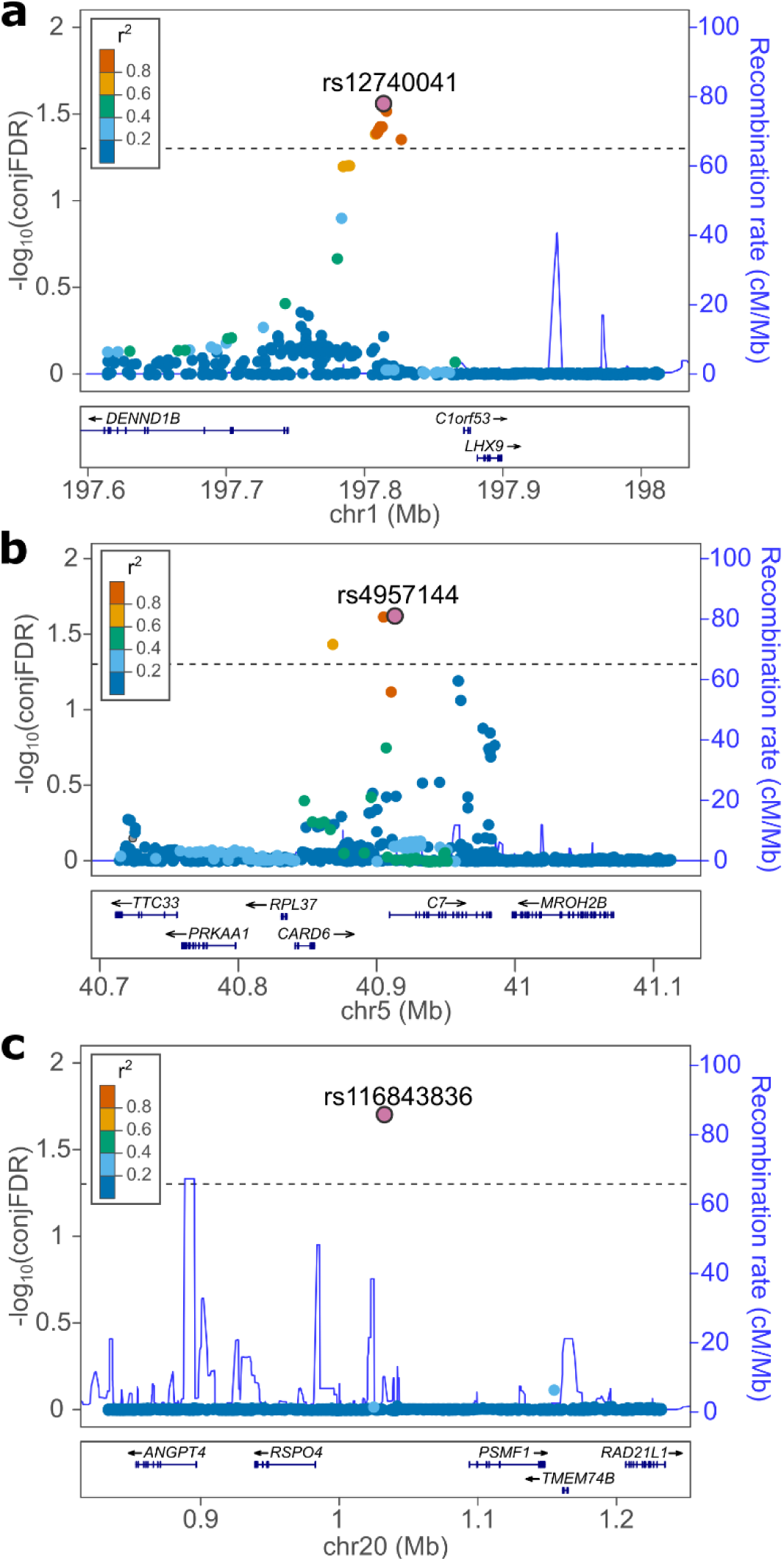
Genetic context of loci identified in the conjFDR analysis. The x-axis represents the positional location of the variants on the chromosome, left y-axis shows -log_10_(conjFDR) value if the variants. In each subplot, a variant with the strongest association is shown in the large purple circle. The color of the remaining markers reflects the degree of linkage disequilibrium (LD), with the strongest-associated variant measured as r^2^ coefficient (described in the legend). The recombination rate is plotted as a solid blue line; its value in centimorgan/megabase (cM/Mb) is indicated on the right y-axis. The black dashed lines indicate the conjFDR threshold = 0.05. Surrounding of the loci with the strongest signal at: **a**, rs12740041 (conjFDR=2.73E-02); **b**, rs4957144 (conjFDR=2.37E-02); **c**, rs116843836 (conjFDR=1.96E-02).

### Identification of relevant tissues and cell types

A total of 7 datasets with tissue/cell type-specific gene expression and chromatin state data were assessed to determine potential enrichment of tissue/cell type-specific categories in MSA heritability. Among all datasets, we identified elevated enrichment for blood/immune and digestive categories in the Roadmap dataset [22] (Supplementary Figure 2a right, and Supplementary Table 2). Furthermore, the ImmGen dataset [23] highlighted a trend for enrichment for B and Myeloid cells in MSA (Supplementary Figure 2, b, left and Supplementary Table 2). Taken together, these results suggested the relevance of the immune and digestive system in MSA pathogenesis. Though we observed a trend for enrichment for various categories, they were not significant when corrected for multiple testing, most likely due to the small sample size of our MSA cohort (Supplementary Figure 2, Supplementary Table 2).

### Functional annotation of identified loci

Functional annotation of three leading variants from loci shared between MSA and IBD/CD as identified in conjFDR analysis (conjFDR < 0.05) showed that one variant (rs4957144) is located in the first intron of *C7* gene while two others: rs12740041 and rs116843836 are intergenic variants located upstream *DENND1B* and *RSPO4* genes correspondingly (Table 1). A lead variant of locus shared between MSA and IBD/CD on chromosome 5 at 5p13.1 (rs4957144, CADD = 14.2) has a CADD score above 12.37 suggesting deleteriousness [24] (Supplementary Table 1). Querying these three variants for eQTL status in the GTEx data (version 7) [25] revealed *DENND1B* (rs12740041), *TTC33* (rs4957144) and *PSMF1* (rs116843836) as potential target genes in various tissues (Table 2). Additional scan of blood (N > 30,000) [26] and brain (N > 520) [27] eQTL summary statistics with substantially larger sample sizes than corresponding tissues in GTEx (N_blood_<400, N_brain_<200) suggested rs12740041 as eQTL for *DENND1B* and rs4957144 as eQTL from *TTC33* in both blood and brain, while rs4957144 was also highlighted as potential eQTL for *RPL37* gene in brain (Table 2). *TTC33* is 158 kb downstream of rs4957144 (Supplementary Figure 3), thus is not shown in Table 1, however, we decided to include it into consequent analyses together with genes listed in Table 1. Assessing a single-variant of eQTL data is prone to false positives [28]. We applied LocusCompare (see URLs) to refine our eQTL findings and check whether loci identified in conjFDR analysis colocalizes with eQTL signal [29]. Indeed, a notable colocalization was observed for rs12740041, where a clump of variants in LD (r^2^>0.4) significantly associated with MSA, CD, and IBD in the conjFDR analysis also revealed substantial deregulation of *DENND1B* gene expression in the brain (Figure 5). The same locus also revealed pronounced colocalization of conjFDR and eQTL signals in other tissues (Supplementary Figure 4, b, c). Observed colocalization of conjFDR and eQTL signals in multiple tissues suggests that deregulation of *DENND1B* gene expression is more likely to be involved in MSA/IBD/CD pathogenesis. Other loci identified in our analysis did not show strong evidence of colocalization events (Supplementary Figure 4, d - j).

**Table 2.** Significant eQTL functionality of lead variants of loci shared between MSA and IBD/CD. Only results passing multiple testing correction from GTEx (passing FDR<0.05), Qi et al. [27] (passing Bonferroni correction p<0.05/N, where N is a number of genes tested for a given SNP) and Võsa et al. [26] (passing Bonferroni correction p<0.05/N). The effect size was computed in a normalized space; thus, its magnitude does not have direct biological interpretation.

| SNP | Gene | Tissue | Data source | p-value | Effect size |
| --- | --- | --- | --- | --- | --- |
| rs12740041 | *DENND1B* | Brain | Qi et al. | 4.88E-04 | 0.18 |
|  |  | Blood | Vösa et al. | 6.90E-25 | 0.12 |
|  |  | Nerve – Tibial | GTE<sub>x</sub> | 6.00E-05 | 0.15 |
| rs4957144 | *TTC33* | Brain | Qi et al. | 3.48E-03 | -0.14 |
|  |  | Blood | Vösa et al. | 1.09E-16 | 0.09 |
|  |  | Artery – Tibial | GTE<sub>x</sub> | 7.50E-07 | 0.13 |
|  | *RPL37* | Brain | Qi et al. | 6.32E-08 | -0.24 |
| rs116843836 | *PSMF1* | Cells - Transformed fibroblasts | GTE<sub>x</sub> | 3.00E-07 | 0.66 |
|  |  | Muscle – Skeletal | GTE<sub>x</sub> | 3.40E-07 | 0.48 |
|  |  | Nerve – Tibial | GTE<sub>x</sub> | 1.30E-06 | 0.44 |

**Figure 5.**
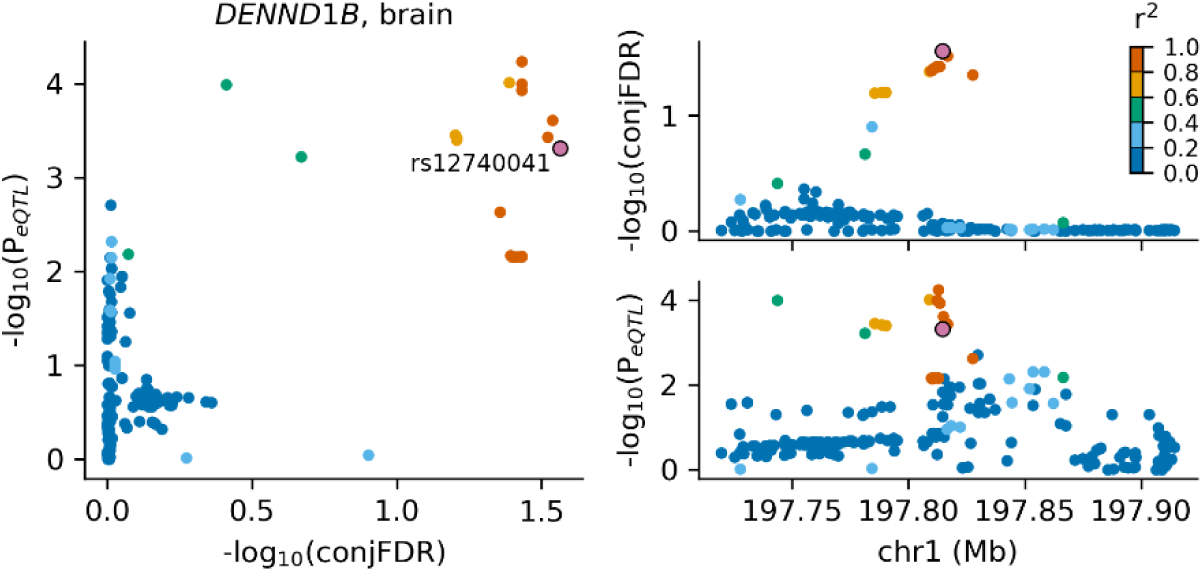
Colocalization of association signals from conjFDR analysis and brain eQTL data [27] in the upstream region of *DENND1B* gene. Scatter plot of the -log_10_(conjFDR) plotted against -log_10_(P_eQTL_) (left), regional association plot for - log_10_(conjFDR) values (top right), regional association plot for -log_10_(P_eQTL_) (bottom right). A lead variant of the locus identified in the conjFDR analysis (rs12740041) is shown in the large purple circle. Other variants are colored according to the degree of linkage disequilibrium (LD) with rs12740041 measured as r^2^ coefficient (described in the legend).

### Gene-level association analysis

Genes within 100kb from lead SNPs identified in the conjFDR analysis were assessed to estimate the cumulative impact of coding variants observed in IBD/CD and MSA cohorts. SKAT test [30] applied to all coding variants (no minor allele frequency threshold) identified *C7* gene as a top association in IBD, CD, and MSA (p_MSA_=1.10E-03, p_CD_=2.74E-05, p_IBD_=5.99E-05) suggesting the importance of the genetic variability within *C7* gene for MSA and IBD/CD (Table 3). Three other genes from the same locus: *CARD6* (p_IBD_=3.33E-02), *RPL37* (p_MSA_=8.79E-03) and *TTC33* (p_CD_=9.46E-03) were nominally significant in IBD, MSA, and CD correspondingly but did not survive multiple testing correction.

**Table 3.** SKAT test p-values in MSA, CD and IBD cohorts for genes within 100 kb from leading SNPs identified in the conjFDR analysis of MSA vs IBD/CD at conjFDR<0.05 (Table 1) and eQTL genes of these leading SNPs (Table 2). p-values for all three cohorts (MSA, CD, and IBD) are estimated using coding variants with no minor allele frequency threshold. Nominally significant p-values (p-value < 0.05) are typed in *italic*, p-values surviving Bonferroni correction (p-value < 0.05/9 = 5.55E-3) are typed in **bold**. “-”: data not available because the gene was not screened in the corresponding cohort.

| Lead SNP | Genes within 100 kb | p-value in SKAT test |  |  |
| --- | --- | --- | --- | --- |
|  |  | MSA | CD | IBD |
| rs12740041 | *C1orf53* | 9.95E-01 | 6.50E-01 | 7.02E-01 |
|  | *LHX9* | 4.60E-01 | 7.86E-01 | 9.24E-01 |
|  | *DENND1B* | 8.21E-01 | - | 9.42E-01 |
| rs4957144 | *C7* | **1.10E-03** | **2.74E-05** | **5.99E-05** |
|  | *CARD6* | 5.38E-01 | 4.12E-01 | 3.33E-02 |
|  | *RPL37* | 8.79E-03 | - | - |
|  | *MROH2B* | - | - | - |
|  | *TTC33* | 7.67E-01 | 9.46E-03 | 4.36E-01 |
| rs116843836 | *RSPO4* | 8.97E-01 | 8.17E-01 | 4.46E-01 |
|  | *PSMF1* | 8.36E-01 | 2.79E-01 | 7.05E-01 |

### Gene set enrichment analysis

FUMA gene set enrichment analysis revealed significant overrepresentation of genes located within 100kb from conjFDR lead SNPs in several biological processes (based on Gene Ontology classification) (Supplementary Table 3). Many of these genes were related to immune responses. Complementary to FUMA analysis, our pathway analysis using DEPICT showed that ITGA2 subnetwork was the most significant gene set for lead SNPs shared between MSA and IBD/CD (nominal p=5.55E-07). However, it did not survive multiple testing correction at FDR<0.05. Complete results of this analysis are presented in Supplementary Table 4.

### Dysregulation of candidate genes in the degenerating midbrain of MSA mice

The PLP-hαSyn transgenic model is characterized by MSA-P-like striatonigral degeneration triggered by human alpha-synuclein overexpression in oligodendrocytes and partly mediated by neuroinflammatory responses [31]. We assessed the expression of nine top candidate genes as identified in the conjFDR analysis. *MROH2B (p=0.03)* and *C7 (p=0.04)* showed significant dysregulation in the midbrain of MSA mice following multiple t-test comparisons corrected with the Bonferroni-Dunn method (Figure 6, Supplementary Table 5). However, the expression levels of *MROH2B* gene were negligible, therefore we conclude that among the examined genes *C7* is the top candidate linked to MSA-like neurodegeneration in the transgenic mouse model.

**Figure 6.**
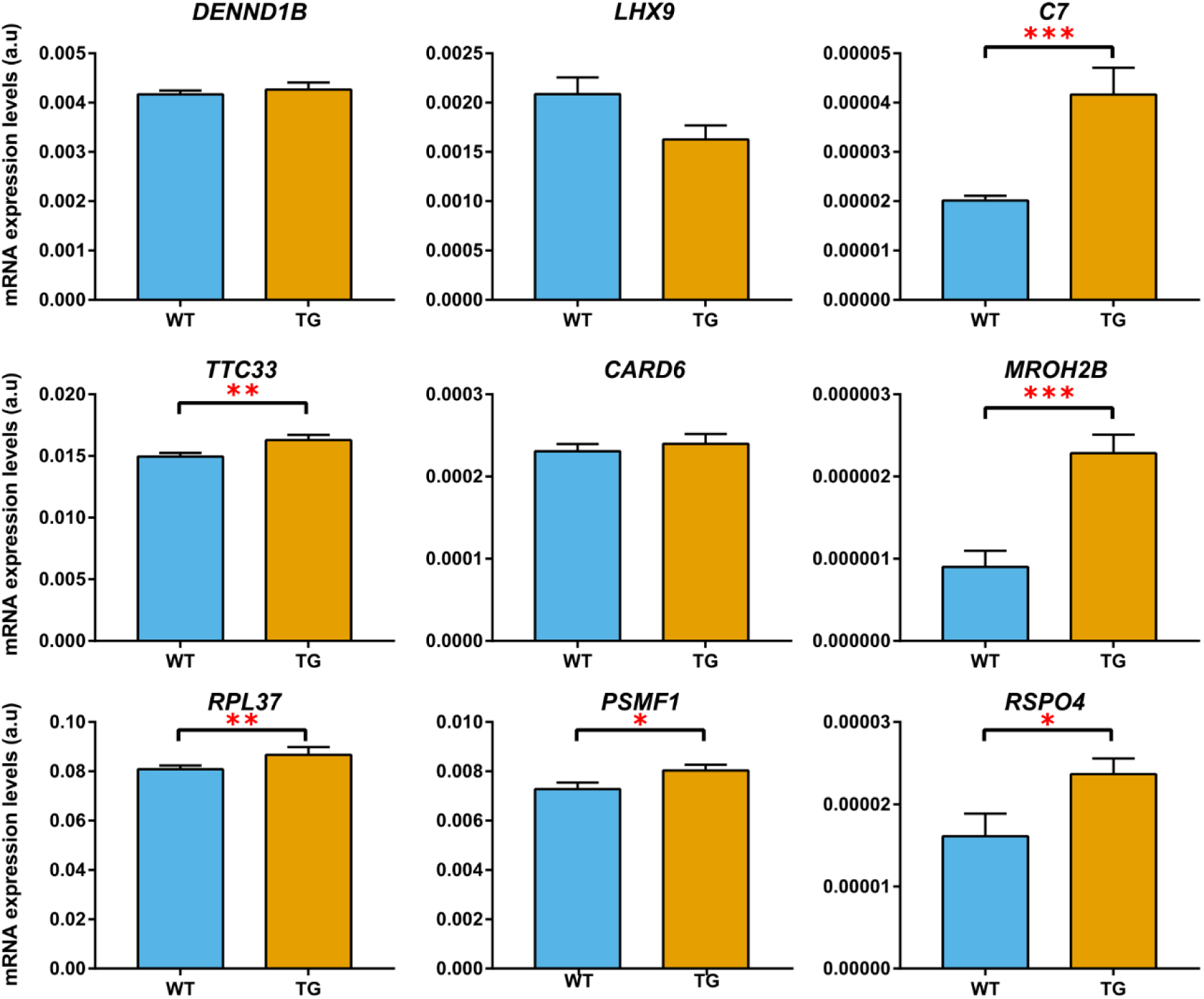
Comparison of expression levels in the midbrain between wild-type mice (blue bars) and transgenic MSA mice (orange bars) for genes identified in the conjFDR analysis. Genes with significantly different expression between wild-type and transgenic mice are marked either with * (nominal p-value < 0.05), with ** (nominal p-value < 0.01) or with *** (nominal p-value < 0.005). *C7* and *MROH2B* genes survive multiple testing with Bonferroni-Dunn correction (adjusted p< 0.05). Error bars show standard error of the mean.

## DISCUSSION

The genetic studies so far failed to provide disease-associated genetic loci for MSA [17, 32]. Nevertheless, because of clinical overlap with PD, a number of studies have assessed the association of top genetic loci of PD with MSA [33, 34]. Thus far, results on the underlying genetic heterogeneity in MSA pathogenesis have been conflicting [32]. The present study, using a genome-wide genetic pleiotropy informed approach, provided evidence of shared genetic etiology between IBD/CD and MSA.

The use of a genetic-pleiotropy informed approach has been successful in identifying shared loci for many complex diseases, including AD and PD [35]. Indeed, a previously published study showed a substantial genetic overlap between PD and autoimmune diseases, in particular with CD and T1D [6]. These studies highlight the involvement of the immune component in neurodegenerative diseases. Given that MSA is a synucleinopathy, our study followed an unbiased approach to understand the extent of the shared genetic etiology between MSA and autoimmune diseases. Q-Q plots (Figure 2) suggested a genetic overlap between MSA and IBD/CD. Our approach helped to expand the genetic spectrum of MSA pathogenesis. These putative loci have not been reported in the previously published MSA GWAS and highlight the utility of such agnostic approaches in gene discovery [32].

MSA GWAS data have yielded negligible heritability estimates in a range of 2% to 6% [17]. The lack of identifying heritability in MSA could be due to studies not using a tissue and cell-specific data in conjunction with GWAS data, which has been shown to increase power in deciphering the missing heritability [36]. Furthermore, as it has been shown in various other complex diseases, including neurodegenerative diseases, simultaneous inclusion of common and rare variants increases power to detect the underlying genetic underpinnings for complex diseases [37-40]. By leveraging various gene expression datasets, our study showed that MSA heritability is enriched in genes with the highest specific expression in blood and digestive categories in GTEx and Roadmap datasets. Interestingly, we observed evidence of enriched heritability in genes showing elevated expression of immune cells in our study. Though the elevated expression was not statistically significant, this is likely due to the small sample of our MSA GWAS data [32]. Our FUMA analysis provided further evidence of involvement of a locus on chromosome 5 at 5p13.1, which encompasses the *C7* gene and shows a high deleterious and moderate regulome score, indicating genetic variability within *C7* gene could modulate disease risk. The role of the *C7* gene was further strengthened by detecting a significant association of common and rare variants of *C7* in both phenotypes, indicating that genetic variability within *C7* gene is important for disease pathogenesis. This genetic evidence is further supported by the findings in the PLP-hαSyn transgenic model of MSA-P-like neurodegeneration, which showed a significant difference in the expression of *C7* gene in the midbrain, a region specifically characterized by early neuroinflammatory response and neuronal loss linked to oligodendroglial alpha-synucleinopathy [31]. All these converging genetic/functional data provide compelling evidence regarding the *C7* gene as a novel locus for MSA pathogenesis. Nevertheless, further functional studies are warranted to decipher the role of *C7* gene in MSA.

Our study reinforces the potential role of “gut-brain axis” in neurodegenerative diseases, and in particular parkinsonian syndromes [41]. For example, a previously published study showed considerable genetic overlap between PD, schizophrenia, and CD [42]. Interestingly, some comorbidity between PD and CD was driven by the genetic variability within *LRRK2* [43]. Furthermore, a recently published study further refined the *LRRK2* locus signal in CD and identified the p.*N2081D* variant driving increased CD risk [43]. Of note, both p.*G2019S* and p.*N2081D* mutations are located in the kinase domain of *LRRK2*, highlighting the role of kinase activity in disease pathogenesis [43]. *LRRK2* has been shown to be involved in a diverse range of functions, including vesicular trafficking, autophagy, and inflammation [44].

A recently published study using population-based data from Denmark showed a 22% increased risk for PD for patients with IBD as compared to non-IBD individuals. Interestingly, the authors also observed a trend for increased risk for MSA in IBD patients [45]. This indicates a bi-directional mechanism between the central and enteric nervous system mediated via the gut microbiome, which is directly linked to the status of the intestinal immune system [41]. Thus, the chronic state of inflammation can promote systemic inflammation and neuroinflammation [46]. Interestingly, a recently published study showed the migration of human α-synuclein after injecting into the intestinal wall of rats to the dorsal motor nucleus in the brainstem in a time-dependent manner, suggesting that changes induced in α-synuclein in the gut can directly affect the brain and elicit an immune response [47].

The loci identified in our study are directly linked to immune-related activities. A recently published study identified genetic variability in *C7* gene as a major risk factor for AD in the Han Chinese population [48]. In addition, the complementary system has been shown to be implicated in a diverse range of functions, including Aβ clearance, microglia activation, neuroinflammation, apoptosis and neuron death [12, 49]. Of note, in our epigenetic analyses, we obtained significant evidence of involvement of *DENND1B* (DENN Domain Containing 1B) gene in blood. *DENND1B*, a member of the connecdenn family, has been shown to play a role in clathrin-mediated endocytosis [50]. The evidence obtained from our study underlying the relevance of immune and vesicular trafficking is important for MSA pathogenesis. Our pathway analysis identified ITGA2 pathway as a “top hit”. Integrins are cell adhesion mediators and have been shown to be involved in a diverse range of human diseases, including IBD, as has been shown in a recently published GWAS [51].

Neuroinflammation characterized by microglia activation has been involved in neurodegenerative diseases, including MSA [46, 52]. Thus, interventional strategies targeting the neuroinflammation offers an alternative approach to either halt or slow disease progression. For example, a study by Mandler et al. [47] performed active immunization againts α-synuclein in MBP α-SNCA transgenic mice. They observed reduced α-synuclein colocalization in oligodendrocytes and astrocytes along with reduced neuronal death and motor deficits. Similarly, therapeutic strategies, which interfere with the synthesis of TNF-α, have been tested in animal models of PD [47], though results generated so far have been remained inconclusive.

The current results might have potential clinical implications. The more extensive clinical evaluation of patients with IBD/CD for monitoring immune/inflammatory and MSA related symptoms should be allowed, as has been suggested in a recently published study [45].

In conclusion, our study extends the genetic architecture of MSA and provides evidence of shared genetic etiology with IBD. Importantly, our findings extend the “gut-brain axis” spectrum from PD to atypical parkinsonian syndromes.

## URLs

condFDR/conjFDR software, https://github.com/precimed/pleiofdr; LocusCompate tool, http://locuscompare.com;

## METHODS

### Participant samples

We used summary statistics from 7 autoimmune disease GWAS and independent GWAS on MSA: IBD (25042 cases, 34915 controls) [51], CD (12194 cases, 34915 controls) [51], ulcerative colitis (12366 cases, 34915 controls) [51], diabetes mellitus type 1 (7514 cases, 9045 controls) [53], celiac disease (4533 cases, 10750 controls) [54], rheumatoid arthritis (29880 cases, 73758 controls) [55], multiple sclerosis (9772 cases, 17376 controls), and MSA (918 cases and 3864 controls) [32]. IBD, CD and ulcerative colitis GWAS [51] were conducted with the same set of controls. IBD cases used in [51] included CD, ulcerative colitis and unclassified cases. Details of the inclusion criteria and phenotype characteristics of the GWAS are described in the original publications. The relevant institutional review boards or ethics committees approved the research protocol of the individual GWAS used in the present analysis, and all participants gave written informed consent.

### Genetic overlap between MSA and autoimmune diseases

To visually assess enrichment of cross-phenotype polygenic overlap between MSA and autoimmune diseases, we generated conditional quantile-quantile (Q-Q) plots by conditioning MSA on each of seven autoimmune phenotypes and vice versa [19]. Q-Q plots are commonly used for visualization of p-values from GWAS summary statistics to assess enrichment of association by plotting quantiles of the observed distribution of association p-values with the phenotype against quantiles of p-value distribution expected under no association (standard uniform distribution). In the absence of association, the Q-Q plot represents a straight line (diagonal of the first quadrant), while deflection from the line indicates the presence of a systematic association. Conditional Q-Q plots are constructed by defining subsets of variants based on significance levels in the conditional phenotype and constructing Q-Q plots for associated-values in the primary phenotype for each subset separately. Enrichment of genetic overlap between primary and conditional phenotype emerges in the conditional Q-Q plot as successive leftward deflections as the significance with the conditional phenotype increases. For more details, please see [19]. Prior to the construction of Q-Q plots, the genomic inflation control procedure described in [19] was applied to correct p-values for all analyzed phenotypes. Additionally, to correct for LD-induced inflation, the contribution of each SNP was measured considering LD structure in the surrounding region, using 100 iterations of random pruning at LD threshold r^2^=0.1. LD structure (r^2^ values) was estimated with PLINK [56] using the 1000 Genomes Project phase 3 European subpopulation data [57]. At each iteration, a set of nearly LD-independent SNPs was selected by taking one random SNP in each LD-independent region (clump of SNPs in LD, r^2^>0.1). The final contribution of each SNP was estimated as an average across all iterations. Because of the underlying complex LD structure encompassing the *HLA* and the *MAPT* regions, which may inflate a conditional Q-Q plot, and introduce a bias in the conjDFR estimation, we excluded SNPs from these regions (hg19 locations chr6:25119106-33854733 and chr17:40000000-47000000, respectively) [58]. The data corrected for LD-induced inflation, including the HLA and *MAPT* regions, were used for the downstream analyses.

### Shared loci between MSA and IBD

The phenotypes which showed substantial genetic overlap with MSA in conditional Q-Q plots were further analyzed with genome-wide conjFDR method to identify shared genetic loci between MSA and autoimmune diseases [19] (see URLs). The conjFDR method is based on the condFDR approach. The condFDR method combines summary statistics from two phenotypes (primary and conditional) and estimates a posterior probability that a variant has no association in the primary phenotype, given that p-values of the variant in both primary and conditional phenotypes are lower than observed p-values. Therefore, condFDR may boost the discovery of loci associated with a primary phenotype by leveraging associations with conditional phenotype. The increase in statistical power is achieved by re-ranking variants as compared to ranking based on the original GWAS p-values [19]. On the contrary, the ranking induced by the standard unconditional FDR (e.g. Benjamini–Hochberg procedure) does not change the order of variants as compared to nominal p-values.

The conjFDR is an extension of condFDR allowing identification of loci associated with both phenotypes simultaneously. For each specific variant, conjFDR is defined as a maximum of two condFDR values (taking one phenotype as primary and another as conditional and then swapping their roles). Therefore, conjFDR provides a conservative estimate of a posterior probability that a variant has no association with either of the phenotypes, given that the p-values for that variant in both analyzed phenotypes are lower than the observed p-values. For more details please refer to the original publication [19].

### Identification of relevant tissues and cell types

To understand whether conjFDR genetic-pleiotropy analyses help further to explain the heritability in MSA by identifying genes expressed in certain tissue/cell types, we used a modified LD score regression applied to specifically expressed genes (LD-SEG) method [36]. In brief, for each gene a t-test was evaluated for the tissue/cell type-specific expression level. Subsequently, the top 10% of genes were selected and a 100 kb window was added around the transcribed region of each selected gene. Finally, stratified LD score regression [59] was applied to resulting genetic regions. This procedure was then repeated for each tissue/cell type of interest. We used publicly available tissue/cell type datasets together with corresponding precomputed gene sets and respective LD-scores as described in the LD-SEG method [36]. A total of seven datasets were used for analyses, including data from GTEx (53 tissues and 13 brain regions) [60], Franke lab (152 tissues) [61, 62], Roadmap Epigenomics consortium (396 tissues and cell types) [22], EN-TEx (93 tissues and cell types), Cahoy et al. [63] (3 brain cell types), ImmGen Consortium (292 immune cell types) [23] and Corces at al. [64] (13 cell types).

### Functional mapping and annotation of identified loci

Two types of analyses were performed. First, positional and functional annotation of lead variants of the loci identified in the conjFDR GWAS analyses was performed using ANNOVAR [65] as implemented in FUMA [66]. Lead variants were annotated with Combined Annotation Dependent Depletion (CADD) [24] scores, measuring the degree of variant deleteriousness on protein structure/function, RegulomeDB [67] scores, predicting the likelihood of regulatory functionality, and chromatin states, estimating transcription/regulatory effects from chromatin states at the locus [22]. Lead variants were queried for known expression quantitative trait loci (eQTLs) in the genotype tissue expression (GTEx) portal [60]. Additionally, eQTL status for lead variants identified in the conjFDR analysis was checked in brain eQTL summary statistics extracted from [27] and blood eQTL summary statistics from [26]. LocusCompare tool (see URLs) was then applied to check whether loci identified in conjFDR analysis colocalizes with eQTL signal.

### Gene-level association tests

The top loci from the conjFDR analysis were further evaluated to determine the role of common and rare variants in IBD/CD and MSA cohorts respectively.

An exome-wide chip covering a total of 205,313 SNPs was used to determine the role of common and rare variants in IBD/CD. RAREMETALWORKER was used with the first ten principal components from PCA as covariates to analyze individual CD and IBD studies and to generate association summary statistics [68]. The exome chip genotype data was converted into Variant Call Format (https://github.com/samtools/hts-specs) format with PLINK [56], and variants were functionally annotated with EPACTS (v3.2.3; http://genome.sph.umich.edu/wiki/EPACTS).

A two-step approach was followed: (i) SKAT test [30] was performed in German CD cohort (exome array study including 4,989 cases and 16,307 controls, using the chip with 205,313 mostly rare coding variants) followed by replication (ii) in a combined IBD cohort (15,236 cases and 34,668 controls). The total number of genes covered in CD cohort was: n = 15,975 for all variants, n = 15,742 for rare variants, and in IBD cohort was: n = 16,306 for all variants, n = 16,138 for rare variants. SKAT test in both cohorts was conducted using all coding variants within identified genes (Table 3). P-values resulting from these tests were corrected for multiple testing using Bonferroni correction.

Whole exome sequencing data from 358 European-ancestry MSA patients and 1,297 neurologically healthy, European controls were generated using the SureSelect Exome target enrichment technology according to the manufacturer’s protocol (Agilent, CA, USA). Sequence alignment to the human reference genome (hg19) and variant calling were performed using the Genome Analysis Toolkit. After removal of duplicate samples using Picard software, stringent quality control filters were applied, including the removal of non-European ancestry individuals and cryptically related individuals, exclusion of individuals with the discrepancy between reported sex and genotypic sex and individuals with high genotype missingness or extreme heterozygosity. Next, all variants were annotated using ANNOVAR [65]. Principle components were generated using flashPCA. SKAT test was performed on all coding variants within identified genes using RVtests and corrected for multiple testing using Bonferroni correction (Table 3).

### Gene set enrichment analysis

DEPICT (depict_140721) [61] was applied to get biological insights from lead variants of loci shared between MSA and overlapping autoimmune diseases, as identified in the conjFDR analysis with relaxed significance threshold (conjFDR<0.35) (Supplementary Table 6). DEPICT is a phenotype-agnostic data-driven integrative method that employs reconstituted gene sets based on massive numbers of experiments measuring gene expression to (1) prioritize genes and pathways related to observed genetic associations and (2) highlight tissues where prioritized genes are highly expressed. Gene set enrichment analysis in DEPICT is based on a ‘guilt-by-association’ procedure [62] which prioritizes genes that share predicted functions with genes from the other associated loci more often than expected by chance. Precomputed GWAS based on arbitrarily generated phenotypes is used to estimate the distribution of p-values under null expectation and to perform multiple testing correction by computing FDR.

In our analysis, we followed an analysis protocol as described in online DEPICT documentation (https://data.broadinstitute.org/mpg/depict/documentation.html). For MSA and overlapping autoimmune disease summary statistic first variants were clumped with PLINK [56] (using flags --clump-p1 1e-5 --clump-kb 500 --clump-r2 0.05). Resulting tag variants were analyzed with the standard DEPICT pipeline using a FDR threshold of 0.05. Leading variants of loci shared between MSA and autoimmune diseases were directly used in standard DEPICT pipeline (no clumping was performed since these variants represent LD-independent regions).

Additionally, gene set enrichment test implemented in FUMA [66] was applied to genes located within 100kb from the lead SNPs of the loci shared between MSA and autoimmune diseases as identified by conjFDR analysis (conjFDR<0.05). The hypergeometric test was used to assess enrichment of input genes in gene sets obtained from MsigDB [69, 70] (which integrates information from KEGG [71], REACTOME [72], Gene Ontology [73] and other sources) taking 19,283 protein-coding genes as a background gene set. Multiple testing correction (per data source) was performed using the Benjamini–Hochberg procedure to control FDR at 5% level.

### Gene expression analysis with real-time PCR in the midbrain of transgenic MSA mice

PLP-hαSyn transgenic mice (also called MSA mice [31, 74]) and wild type controls were kept under temperature-controlled pathogen-free conditions on a light/dark 12 h cycle. All experiments were performed according to the ethical guidelines of the EU (Directive 2010/63/EU for animal experiments) and the Austrian Federal Ministry of Science and Research (permission BMFWF-66.011/0018-WF/v/3b/2015). Twelve-month-old male mice (five PLP-hαSyn and five wild type mice) were perfused intracardially with phosphate buffered saline (PBS, pH 7.4, Sigma) under deep thiopental anesthesia. Brains were extracted, and midbrains were quickly dissected and snap-frozen in liquid nitrogen. Samples were stored at -80°C until further processing. Samples were homogenized in TRIzol reagent (Life technologies) with ULTRA-TURRAX T-8 basic tissueruptor (IKA) and RNA was isolated following the manufacturer’s instructions. RNA samples were retrotranscribed into cDNA by using the High-Capacity cDNA Reverse Transcription Kit (Applied-Biosystems). Analyses were performed in a CFX96 Touch™ Real-Time PCR Detection System (Bio-Rad) using iTaq universal probes supermix (Bio-Rad). *Gapdh* mRNA levels were used as an internal normalization control. All statistical analyses were conducted using the software Graph-Pad Prism 7 (Graphpad Software). The mean ± S.E.M was used to present the results. Comparisons were performed with multiple t-tests with Bonferroni-Dunn correction. An adjusted p-value <0.05 was considered statistically significant.

## Supporting information

A full list of the European Multiple System Atrophy Study Group members

## ACKNOWLEDGMENTS

This work is supported by the grants from Multiple System Atrophy Coalition, USA (to M.S.). This work was further supported by the grants from the German Research Council (DFG/SH 599/6-1 to M.S.), the EU Joint Program-Neurodegenerative diseases (JPND; COURAGE-PD to M.S. and T.G. FKZ 01ED1604), and Michael J Fox Foundation (to M.S.).

This work was supported by the Research Council of Norway (#223273, #225989, #248778) South-East Norway Health Authority (#2016-064, #2017-004), KG Jebsen Stiftelsen (#SKGJ-Med-008). (to A.A.S., O.A.A, O.F., K.S.O., S.B.)

This work was supported in part by the Intramural Research Programs of the National Institute of Neurological Disorders and Stroke (NINDS), and the National Institute on Aging (NIA): project numbers Z01-AG000949, 1ZIA NS003154.

This work was supported by a grant of the Austrian Science Fund (FWF) F4414 (to NS).

This work was supported by the German Federal Ministry of Education and Research (BMBF) within the framework of the e:Med research and funding concept (SysInflame grant 01ZX1606A; GB-XMAP grant 01ZX1709). The project was funded by the Deutsche Forschungsgemeinschaft (DFG, German Research Foundation) under Germany’s Excellence Strategy – EXC 2167-390884018. The project received infrastructure support from the DFG Excellence Cluster No. 306 “Inflammation at Interfaces” and the PopGen Biobank (Kiel, Germany). The KORA research platform (KORA, Cooperative Research in the Region of Augsburg) was initiated and financed by the Helmholtz Zentrum München – German Research Center for Environmental Health, which is funded by BMBF and by the State of Bavaria. Furthermore, KORA research was supported within the Munich Center of Health Sciences (MC Health), Ludwig-Maximilians-Universität, as part of LMU innovativ. SHIP (Study of Health in Pomerania) is part of the Community Medicine Research net (CMR) of the University of Greifswald, Germany, which is funded by BMBF (grants 01ZZ9603, 01ZZ0103 and 01ZZ0403) and the Ministry of Cultural Affairs as well as the Social Ministry of the Federal State of Mecklenburg-West Pomerania, and the network ‘Greifswald Approach to Individualized Medicine (GANI_MED)’ funded by BMBF (grant 03IS2061A). Exome chip data have been supported by BMBF (grant no. 03Z1CN22) and the Federal State of Mecklenburg-West Pomerania. Genotype data of the controls that has been used in this study is available on European Genome-phenome Archive (EGA) under accessions EGAD00010000890 and EGAD00010000234. Genotyping array is available under the accession numbers EGAS00001001232. Genotype data of cases are accessible on EGA under accession EGAD00010001158. Genotype array data is available under accession EGAS00001000924.

Computations were performed on the Abel Cluster, owned by the University of Oslo and Uninett/Sigma2, and operated by the Department for Research Computing at USIT, the University of Oslo IT-department (http://www.hpc.uio.no/).

## AUTHOR CONTRIBUTIONS

A.A.S. performed the analyses, participated in study design, wrote the draft of the paper.

M.S. designed the study, secured funding, and wrote the first draft of the paper.

S.M. performed the analyses, wrote the draft of the paper.

D.E. performed the analyses, wrote the draft of the paper.

M.B.M. performed the analyses, wrote the draft of the paper.

A.H. performed transgenic mice experiments, wrote the draft of the paper.

N.S. performed transgenic mice experiments, wrote the draft of the paper.

All authors: commented and revised the draft.

