## Supplementary Figures for "A genome-wide genetic pleiotropy approach identified shared loci between multiple system atrophy and inflammatory bowel disease"

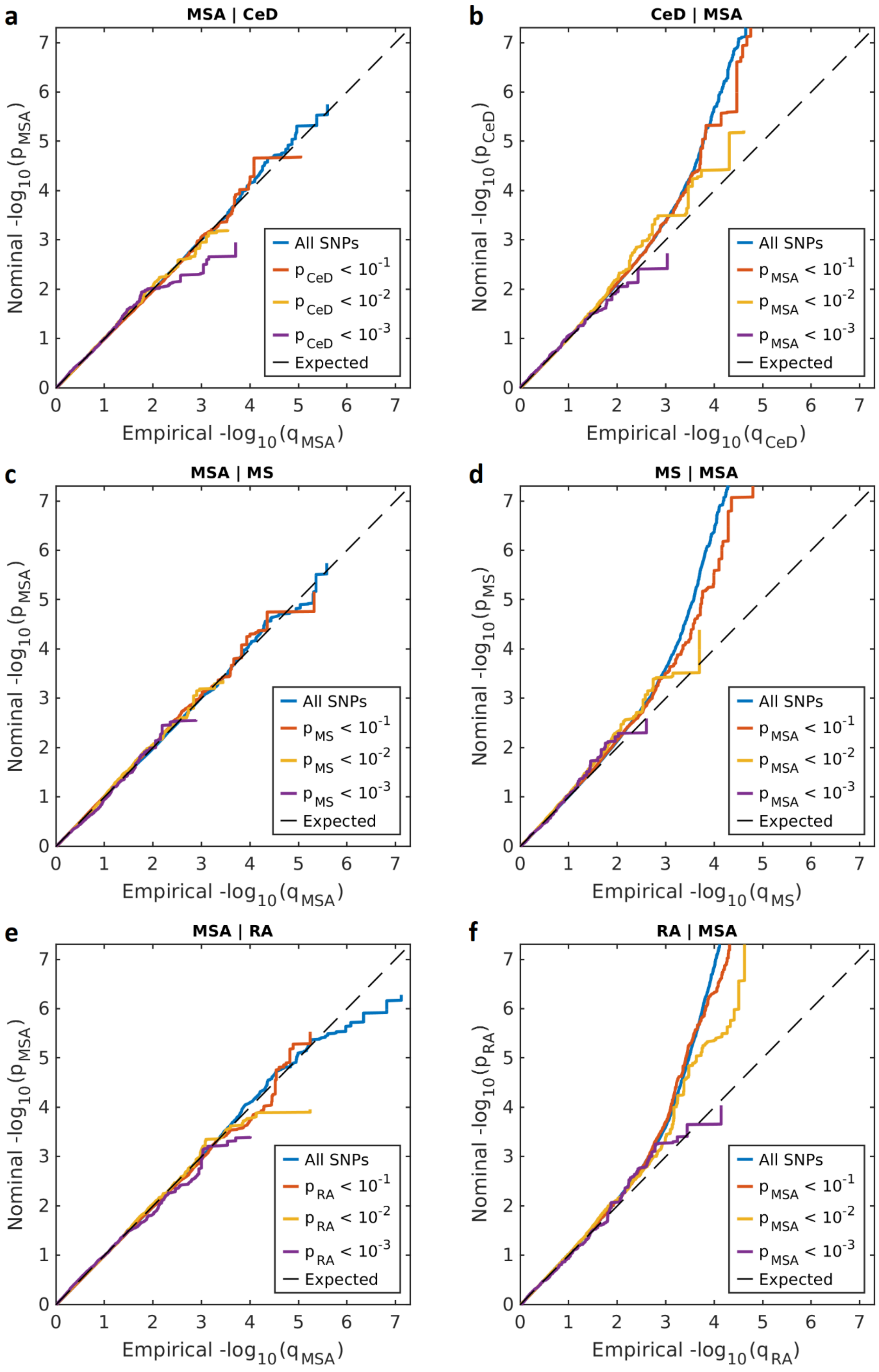

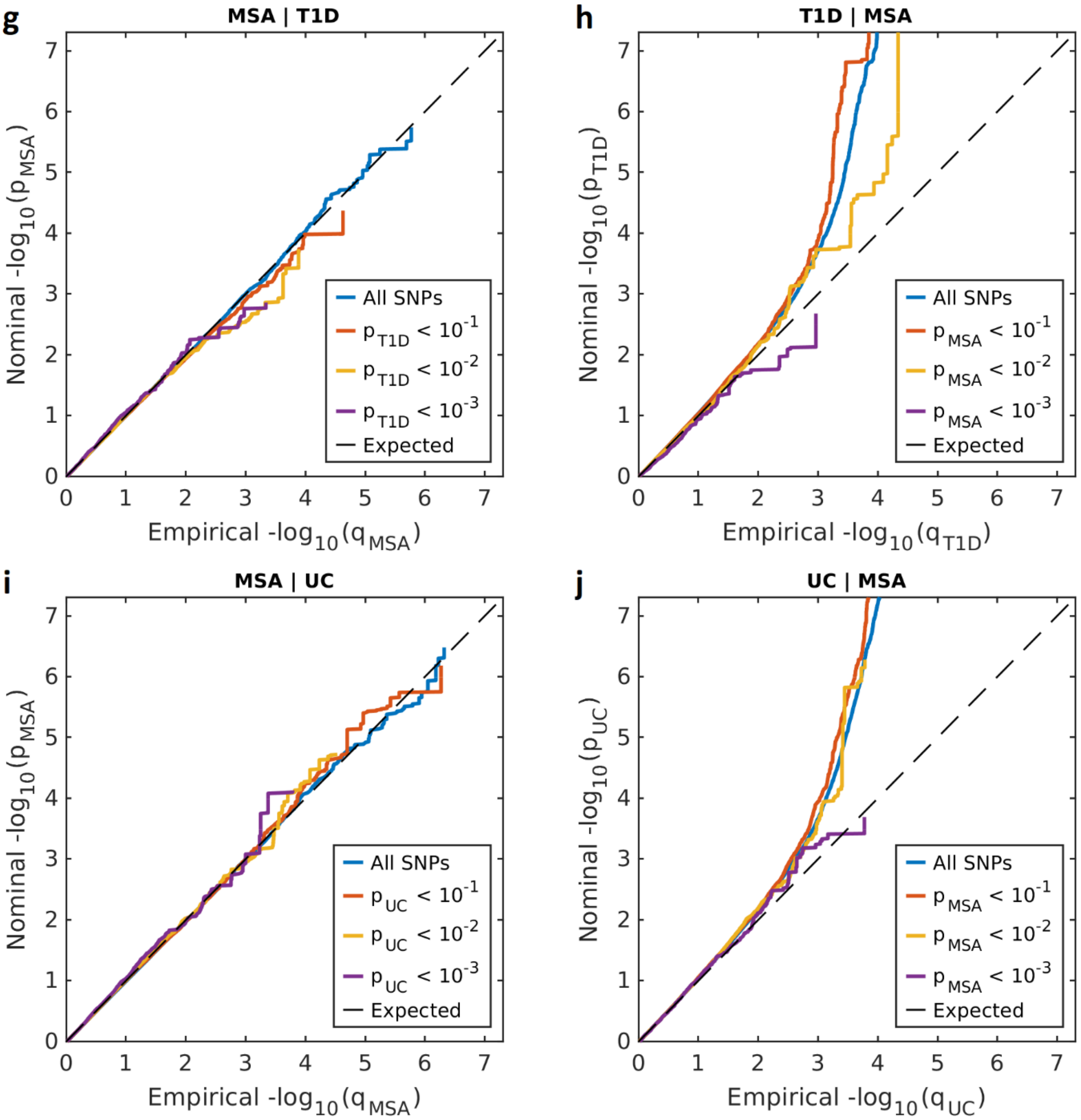


**Supplementary Figure 1**. Conditional Q-Q plot showing show relation between expected (x axis) and observed (y axis) significance of SNPs in the primary phenotype when markers are stratified by their p-values in the conditional phenotype. A sequence of 4 nested strata is presented: all SNPs (blue), $p_{conditional\_phenotype} < 0.1$ (orange), $p_{conditional\_phenotype} < 0.01$ (yellow) and $p_{conditional\_phenotype} < 0.001$ (purple). Dashed black line demonstrates expected behavior under no association. Increasing degree of leftward deflection from the no-association line for strata of SNPs with higher significance in the conditional phenotype indicates polygenic overlap.

A: MSA conditioned on celiac disease (CeD); B: CeD conditioned on MSA; C: MSA conditioned on multiple sclerosis (MS); D: MS conditioned on MSA; E: MSA conditioned on rheumatoid arthritis (RA); F: RA conditioned on MSA; G: MSA conditioned on diabetes mellitus type 1 (T1D); H: T1D conditioned on MSA; I: MSA conditioned on ulcerative colitis (UC); J: UC conditioned on MSA.

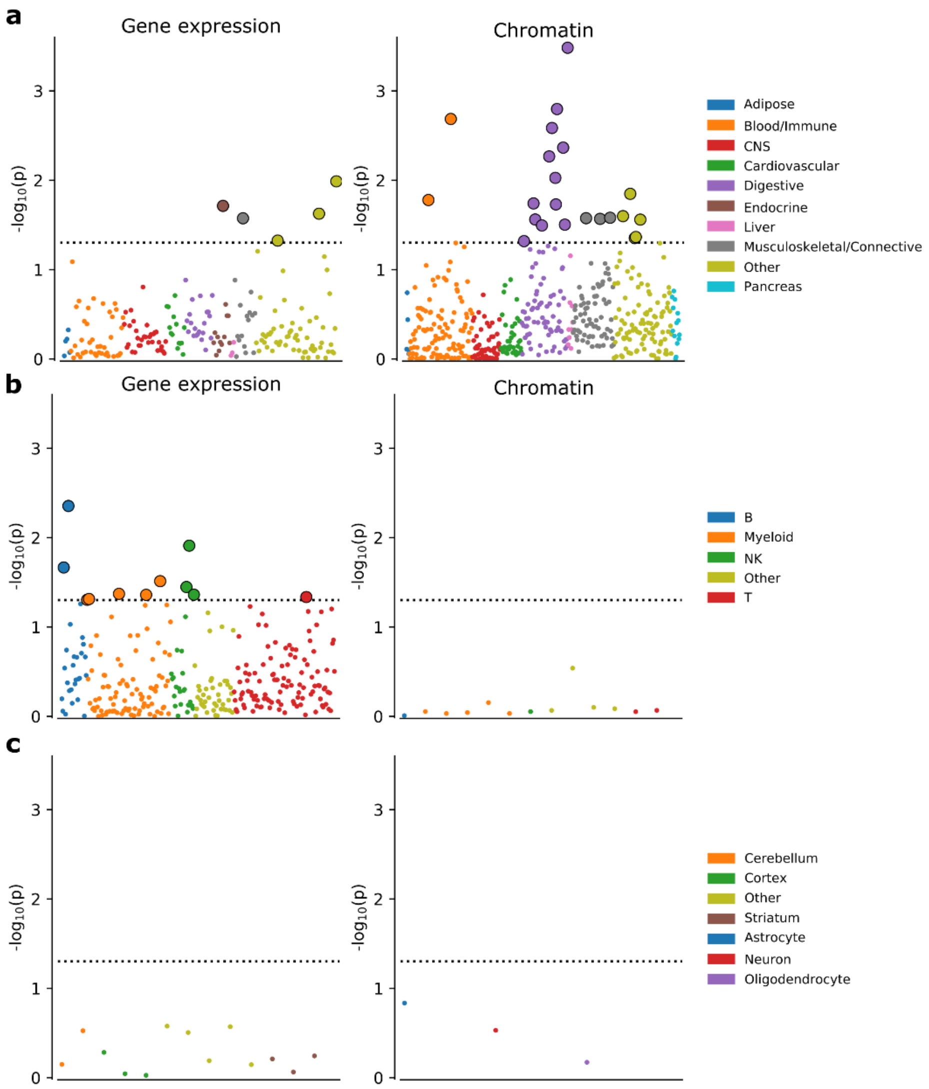


**Supplementary Figure 2.** Tissue/cell-type specific enrichment of MSA heritability. Y-axis shows -log10 uncorrected p-value. None of tissue/cell type is significant after FDR correction with 0.05 threshold. Nominally significant results ($p < 0.05$) are plotted with larger outlined circles. Horizontal dashed line represents nominal significance threshold $-\log_{10}(0.05)$. Numerical results are presented in Supplementary Table 4. **a**, Multiple tissue analysis using gene expression data from GTEx and Franke et al. and chromatin data from Roadmap and EN-TEx datasets (top hit in Digestive category has FDR = 0.16). **b**, Immune cell type analysis using expression data from ImmGen and chromatin data from Corces et al. **c**, Results for different brain regions from GTEx (left), and three brain cell types from Cahoy et al. (right).

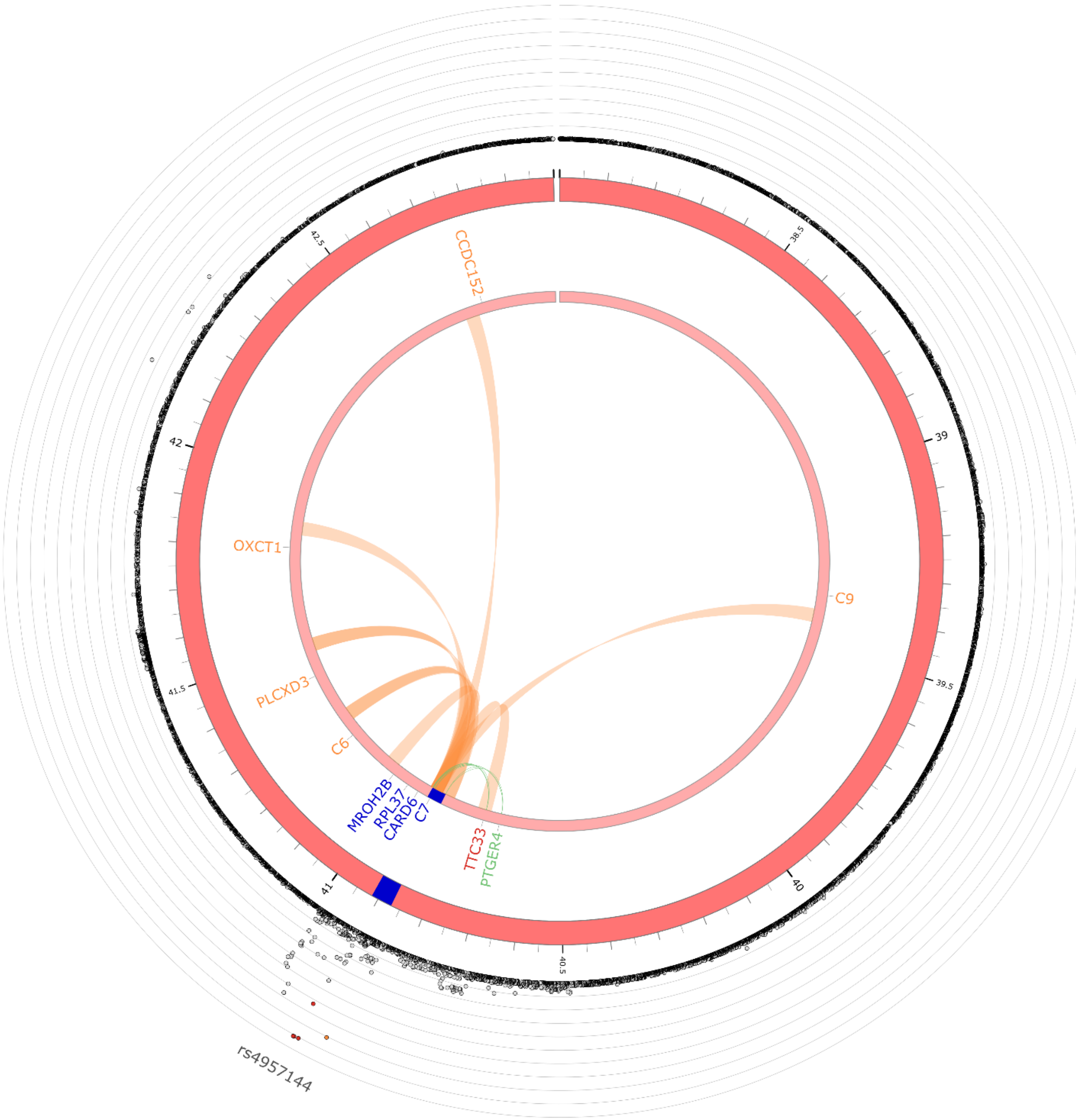


**Supplementary Figure 3.** Circos plot demonstrating locus on chromosome 5 at 5p13.1 identified in the conjFDR analysis (Table 1). Lead SNP is labeled at the outer circle. Genes within 100 kb of the lead SNP have blue labels. Significant eQTL target genes of significant SNPs in the locus have green labels and chromatin interaction regions are showed with orange. The gene has res label if it is both a significant eQTL target and is in the chromatin interaction region.

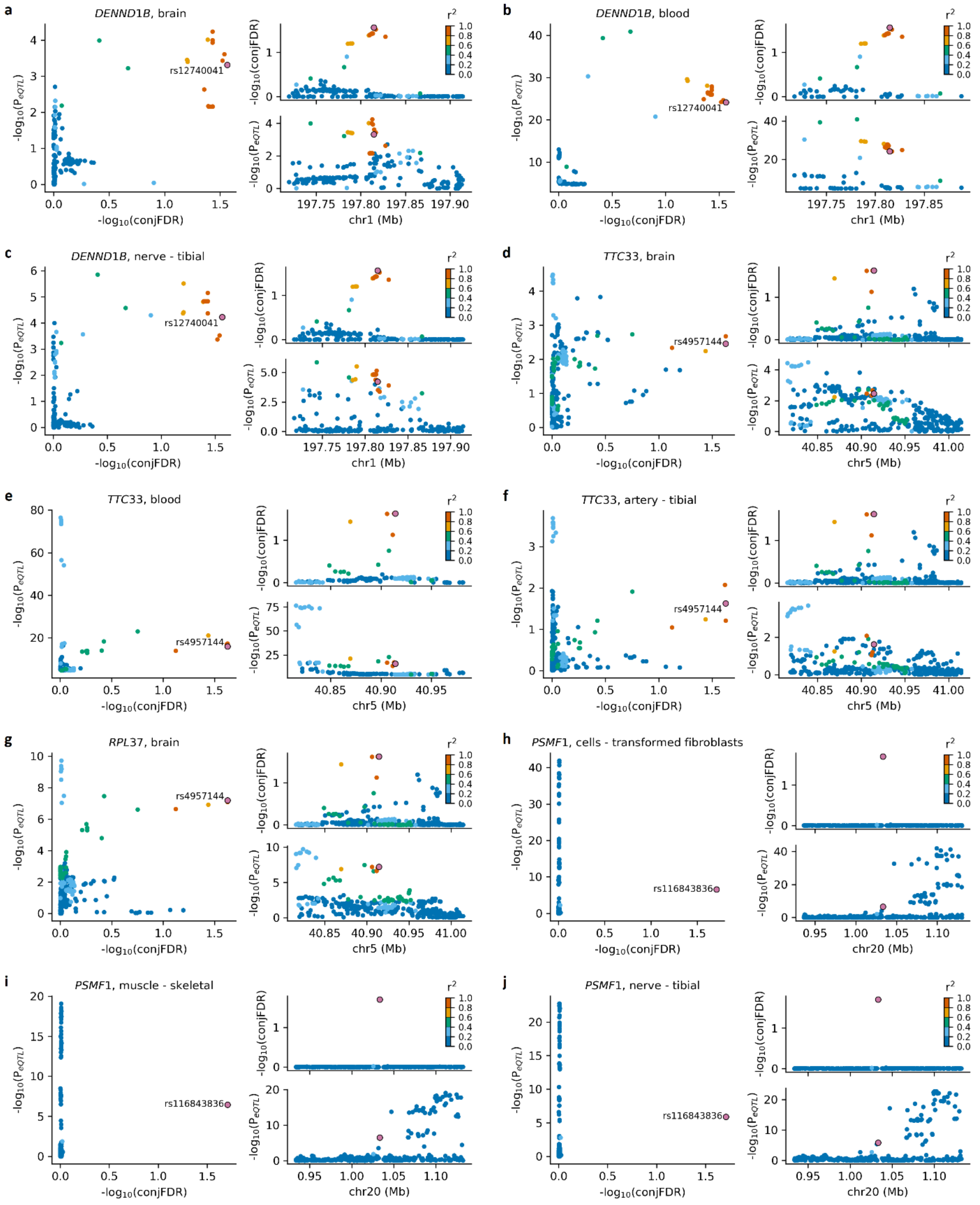


**Supplementary Figure 4.** Colocalization of association signals from conjFDR analysis and eQTL data from multiple tissues for genes within loci identified in conjFDR analysis.

Scatter plot of the $-\log_{10}$(conjFDR) plotted against $-\log_{10}(P_{eQTL})$ (left), regional association plot for $-\log_{10}$(conjFDR) values (top right), regional association plot for $-\log_{10}(P_{eQTL})$ (bottom right). In each subfigure lead variant of the locus identified in conjFDR analysis is shown in the large purple circle and labeled in the scatterplot. Other variants are colored according to the degree of linkage disequilibrium (LD, measured as $r^2$ coefficient) with corresponding lead variant. For each subplot (a - j) the title of the scatterplot shows corresponding gene name and tissue, where brain stands for the eQTL data from [1], blood refers to the eQTL data from [2] and the rest represents various tissues from GTEx.
