## Supplementary material for "A genome-wide genetic pleiotropy approach identified shared loci between multiple system atrophy and inflammatory bowel disease": A full list of the European Multiple System Atrophy Study Group members

**EMSA-SG FULL MEMBERS** (last updated: 2019-07-02)

#1 **Prof. Werner Poewe**, **Prof. Gregor K. Wenning**, University Hospital Innsbruck, Department of Neurology, Anichstrasse 35, **A- 6020 Innsbruck**

#2 **Prof. Karen Østergaard**, Aarhus Kummune- hospital, Department of Neurology, Norrebrogade 44, DK-8000 Aarhus C

#3 **Prof. Wassilios Meissner**, Centre Hospitalier Universitaire de Bordeaux, Service de Neurologie/CHU de Bordeaux, Avenue de Magellan, F-33604 Pessac

#4 **Prof. Olivier Rascol**, **Prof. Anne Pavy-Le Traon**,Toulouse III University, Laboratoire de Pharmacologie, Faculte de Medecine, 37 Allees Jules Guesde, F-31073 Toulouse

#5 **Prof. Thomas Klockgether**, University of Bonn, Department of Neurology, Sigmund-Freud- Straße 25, D-53105 **Bonn**

#6 **Prof. Daniela Berg**, Klinik für Neurologie, Niemannsweg 147, D-24105 Kiel

#7 **Prof. Karla Eggert**, Philipps-University Marburg, Department of Neurology, Rudolf-Bultmann-Straße 8, D-35033 Marburg

#8 **Prof. Thomas Gasser**, Hertie-Institute for Clinical Brain Research, Department of Neurodegenerative Disorders, University of Tübingen, Center of Neurology, Hoppe-Seyler-Str. 3, D-72076 Tübingen

#9 **Prof. Nir Giladi**, **Dr. Tanya Gurevich**, Tel Aviv Sourasky Medical Center, Movement Disorders Unit, Department of Neurology, 6, Weizmann Street, IL-64239 Tel-Aviv

#10 **Prof. Alfredo Berardelli**, Department of Neurology and Psychiatry, Sapienza University Rome, Rome

#11 **Prof. Joaquim Ferreira**, Faculdade de Medicina da Universidade de Lisboa, Instituto de Medicina Molecular, Lisbon

#12 **Prof. Eduardo Tolosa**, Universitat de Barcelona, Hospital Clinic, Department of Neurology, Villarroel, 170, E-08036 Barcelona

#13 **Prof. Håkan Widner**, University of Lund, Department of Clinical Neuroscience, Division of Neurology, Solvegatan 17, SE-22362 Lund

#14 **Prof. Henry Houlden**, University College London, Institute of Neurology, Queen Square, WC1N 3 BG London

#15 **Prof. David J. Brooks**, The Imperial College of Sciences, Technology and Medicine, MRC Cyclotron Unit, 150 DuCane Road,W12 0NN London

#16 **Prof. Andrew Lees**, University of College London, PDS Brain Research Centre, Institute of Neurology, 1 Wakefield Street, WC1N 1 PJ London

#17 **Prof. Ruth Djaldetti**, Department of Neurology, Rabin Medical Cenre, Beilinson Campus, 49100 Petach-Tiqva

#18 **Prof. Alberto Albanese**, Istituto Clinico Humanitas, Via Alessandro Manzoni, 56 I-20089 Rozzano Milano

#19 **Prof. Maria Teresa Pellecchia**, Center for Neurodegenerative Diseases (CEMAND), Department of Medicine and Surgery, Neuroscience Section, University of Salerno

#20 **Prof. Maria Stamelou**, University of Marburg, Germany, Head of Movement Disorders Department, “HYGEIA” Hospital, Athens, Greece, Collaborating consultant and reasearhcer at University of Athens Medical School, 2nd Department of Neurology, Hospital “Attikon”, Athens

#21 **Prof. Vladimir S. Kostić**, University of Belgrade, Institute of Neurology CCS, Ul. Dr Subotića, 11000 Belgrade

#22 **Prof. Pietro Cortelli**, University of Bologna, Department of Neurosciences, Via Ugo Foscolo 30, I-40123 Bologna

#23 **Prof. Heinz Reichmann**, Professor and Chair Dept. Neurology, University of Dresden, Fetscherstrasse 74, D-01307 Dresden

#24 **Prof Tim Lynch**, Dublin Neurological Institute,57 Eccles Street, Dublin 7.

#25 Prof. **Jaroslaw Slawek**, Neurology Dpt. at St. Adalbert Hospital in Gdansk and Dpt. of Neurologic-Psychiatric Nursing at Medical University of Gdansk, Al. Jana Pawła II 50, 80-462 Gdańsk.

#26 **Prof. Bastiaan R. Bloem**, Parkinson Centre Nijmegen (ParC), Department of Neurology, Donders Institute for Brain, Cognition and Behavior, Radboud University Nijmegen Medical Centre, Reinier Postlaan 4, 6500 HB Nijmegen.

#27 **Dr. Alexander Gerhard**, Senior Lecturer in Neurology, Honorary Consultant Neurologist, Greater Manchester Neuroscience Centre, Salford Royal NHS Foundation Trust, Salford, M6 8HD.

#28 **Prof Latchezar Traykov**, **Dr Mariya Petrova**, University Hospital Alexandrovska, 1, Georgi Sofiiski str.,1431 Sofia.

#29 **Prof. Patrick Cras**, Department of Neurology, University of Antwerp, Prinsstraat 13, 2000 Antwerp

**CENTERS (27)**

**Aarhus, Antwerp, Athens, Barcelona, Belgrade, Bologna, Bonn, Bordeaux, Dresden, Dublin, Gdansk, Innsbruck, Kiel, Lisbon, London, Lund, Manchester, Marburg, Milan, Naples, Nijmegen, Petach-Tiqva, Rome, Sofia, Tel-Aviv, Toulouse, Tübingen.**
